# Stimulus Salience Determines the Defensive Behaviors Elicited by Aversively Conditioned Compound Auditory Stimuli

**DOI:** 10.1101/505263

**Authors:** Sarah Hersman, Todd E. Anthony

## Abstract

Animals exhibit distinct patterns of defensive behavior according to their perceived imminence of potential threats. Ethoexperimental [1, 2] and aversive conditioning [3-5] studies indicate that as the probability of directly encountering a threat increases, animals shift from behaviors aimed at avoiding detection (e.g. freezing) to escape (e.g. undirected flight). What are the neural mechanisms responsible for assessing threat imminence and controlling appropriate behavioral responses? Fundamental to addressing these questions has been the development of behavioral paradigms in mice in which well-defined threat-associated sensory stimuli reliably and robustly elicit passive or active defensive responses [6, 7]. In serial compound stimulus (SCS) fear conditioning, repeated pairing of sequentially presented tone (CS1) and white noise (CS2) auditory stimuli with footshock (US) yields learned freezing and flight responses to CS1 and CS2, respectively [6]. Although this white noise-induced transition from freezing to flight would appear to reflect increased perceived imminence due to the US being more temporally proximal to CS2 than CS1, this model has not been directly tested. Surprisingly, we find that audio frequency properties and sound pressure levels, not temporal relationship to the US, determine the defensive behaviors elicited by SCS conditioned auditory stimuli. Notably, auditory threat stimuli that most potently elicit high imminence behaviors include frequencies to which mouse hearing is most sensitive. These results argue that, as with visual threats [8], perceived imminence and resulting intensity of defensive responses scale with the salience of auditory threat stimuli.

## RESULTS

### White noise elicits active fear responses during SCS conditioning irrespective of pairing or stimulus order during training

To define the relationships between CS stimulus properties, their temporal relationship to the US, and the defensive behaviors they elicit, we first asked whether reversing the order of tone (TN) and white noise (WN) presentation during SCS conditioning would also reverse the behaviors these stimuli elicit. To distinguish learned CS-US associations from responses due to sensitization or generalization, an ‘unpaired’ group was also included in which a 60 second gap was introduced between the SCS and the US (**Figures 1A-1F**). As evident from the motion traces (**Figure 1D**), all groups exhibited significantly greater motion during WN than TN, irrespective of the order that these stimuli were presented during training (**Figures 1J-1L**). As conditioning progressed, mice in all groups began to exhibit active responses to the WN, including darting and jumping; these behaviors were quantified using an ‘escape score’ (**Figures 1M-1O**, see Methods) and can be quantitatively distinguished from other behaviors observed during conditioning (e.g. crossings, rapid movements in place; **Figure S1** and **Supplementary Videos**).

**Figure 1.**
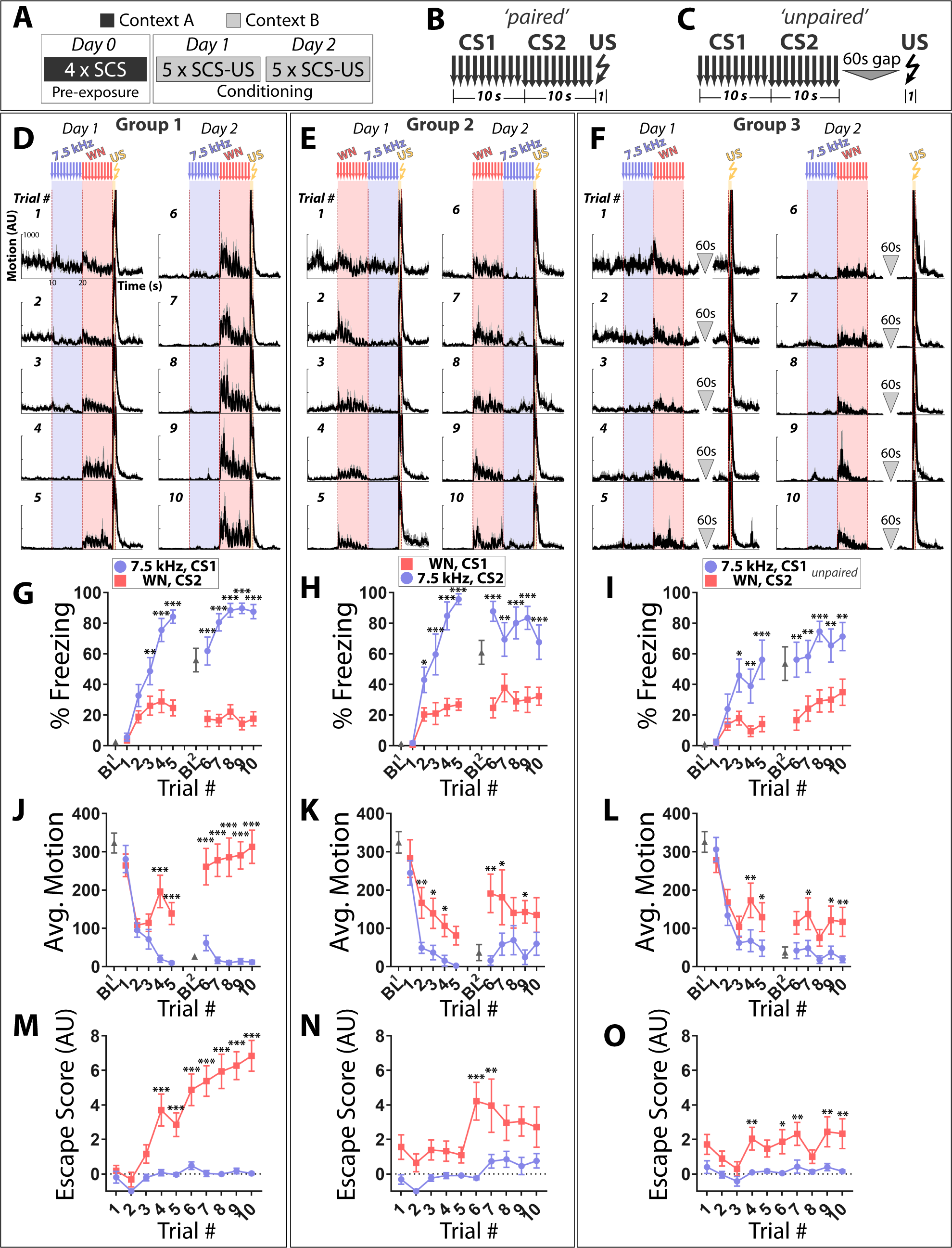
White noise elicits active fear responses during SCS conditioning regardless of temporal relationship to the US. (A-C) Protocol and structure of stimuli applied during conditioning for paired and unpaired groups. (D-F) Motion indices (mean±SEM) showing movement in the absence or presence of stimuli (10x 7.5 kHz pips, 75 dB, 0.5s each at 1 Hz, blue; 10x white noise (WN) pips, 75 dB, 0.5s each at 1 Hz, pink; 1x 0.9 mA footshock, 1 s, yellow) for all 10 conditioning trials (Days 1 and 2). (G-I) Percentage time spent freezing during baseline (BL, 3 minutes prior to the first stimulus presented each day) and trials across each conditioning day. (J-L) Average motion during BL and trials. (M-O) Active fear behavior during each trial quantified as an escape score (see Methods). CS order and pairing: group 1 (D,G,J,M: CS1=7.5 kHz, CS2=WN; *n*=15), group 2 (E,H,K,N: CS1=WN, CS2=7.5 kHz; *n*=10), and group 3 (F,I,L,O: CS1=7.5 kHz, CS2=WN, unpaired; *n*=10). Asterisks indicate significant difference between stimuli for a given trial. Error bars indicate the SEM.

Two observations are additionally noteworthy. First, WN triggered escape behaviors in the unpaired group, although significantly less than in the paired group conditioned using the same SCS (group 1 vs. group 3, F (1, 23) = 19.34, p<0.001), Sidak’s test, significantly different on Trials 8 (p<0.001), 9 (p<0.05) and 10 (p<0.01)). Second, group 1 motion responses to WN on day 2 (**Figure 1D**) were largest immediately following stimulus onset and decreased thereafter until US exposure (paired t-test, average motion first two vs. last two seconds of CS2, trials 6, 7, p<0.01; trials 8, 9, p<0.05). Taken together, these data suggest that imminence in the SCS paradigm is not determined by a cognitive process that calculates time remaining before the impending US, but rather may be related to salience of the auditory stimuli themselves.

### 7.5 kHz tone stimuli promote conditioned flight when presented at high sound pressure levels

Mice can hear sounds from 1 kHz to 100 kHz, but sensitivity to specific frequencies varies dramatically over this range. For example, the minimal sound pressure levels (SPL) that mice can reliably detect for 16 kHz tones is ∼10x lower (10 dB) than for 7.5 kHz tones (20 dB) [9]. Given that the WN stimulus used here and previously [6] is composed of frequencies between 1-20 kHz, one explanation for the above results is that mice hear WN stimuli better than pure 7.5 kHz tones, and so perceive the two stimuli as reflecting distinct points along the threat imminence continuum. One prediction of this model is that a 7.5 kHz CS presented at high SPL should be perceived as more imminent and elicit more escape than the exact same CS presented at low SPL.

To test this prediction, we performed a ‘SPL step test’ in which conditioned mice were presented with a SCS composed of two 7.5 kHz tones: CS1 is held constant at 75 dB while CS2 SPL magnitude begins at 55 dB and is stepped up by 5 dB each trial, finishing at 105 dB (**Figures 2A-2C**). While predominantly freezing was observed at ≤85 dB, the 7.5 kHz SCS began to elicit escape behaviors in the paired group when CS2≥90 dB (**Figure 2H**). Further, escape scores for trials where CS2≥90 dB were significantly higher in group 1 (paired) than group 3 (unpaired): 2-Way Repeated Measures ANOVA, Main Effect of Trial (F (4, 92) = 3.208, p<0.05), Main Effect of Group (F (1, 23) = 4.613, p<0.05. This argues that group 1 responses are influenced by perceived threat levels and are not a simple reflexive reaction to loud sounds. Moreover, escape at later trials was observed in response to CS2 but not CS1, demonstrating that these behavioral changes were not due solely to enhanced responsivity to any stimulus following repeated US exposure.

**Figure 2.**
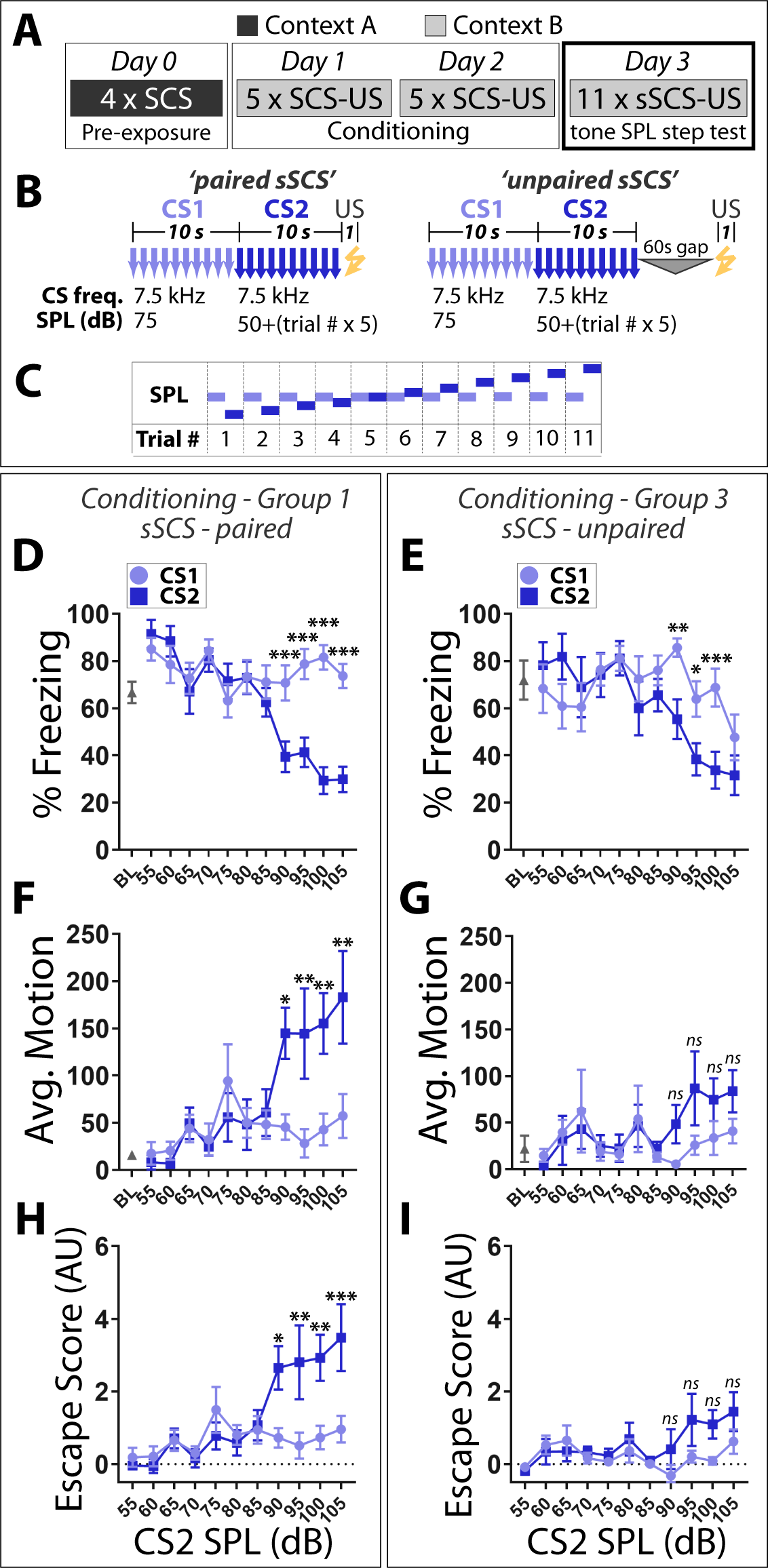
Sound pressure levels determine whether SCS fear conditioned tone stimuli elicit freezing or flight. (A) Mice conditioned in groups 1 and 3 (Figure 1) were run through a tone SPL step test on day 4. (B) The tone step SCS (sSCS) is composed of two 7.5 kHz tone stimuli in which CS1 is held constant at 75 dB while CS2 begins at 55 dB and is stepped up by 5 dB each trial. (C) Schematic of tone SPL step test. (D,E) Percentage time spent freezing. (F,G) Average motion. (H,I) Escape score. Paired sSCS (D,F,H; *n*=15); unpaired sSCS (E,G,I; *n*=10). Asterisks indicate significant difference between stimuli for a given trial. *ns*, not significant. Error bars indicate the SEM.

To determine whether behavioral responses to WN also scale with SPL, we performed a SPL step test using a simple WN CS presented in a novel context (**Figures 3A-3C**). At low SPL (40-45 dB), WN elicited robust freezing and little to no escape behavior. In contrast, at higher SPL (≥60 dB), escape responses were common and freezing was minimal during WN presentations (**Figures 3D-3F**). Thus, SCS fear conditioned 7.5 kHz and WN stimuli elicit freezing or flight behavior according to the SPL magnitude at which they are presented.

**Figure 3.**
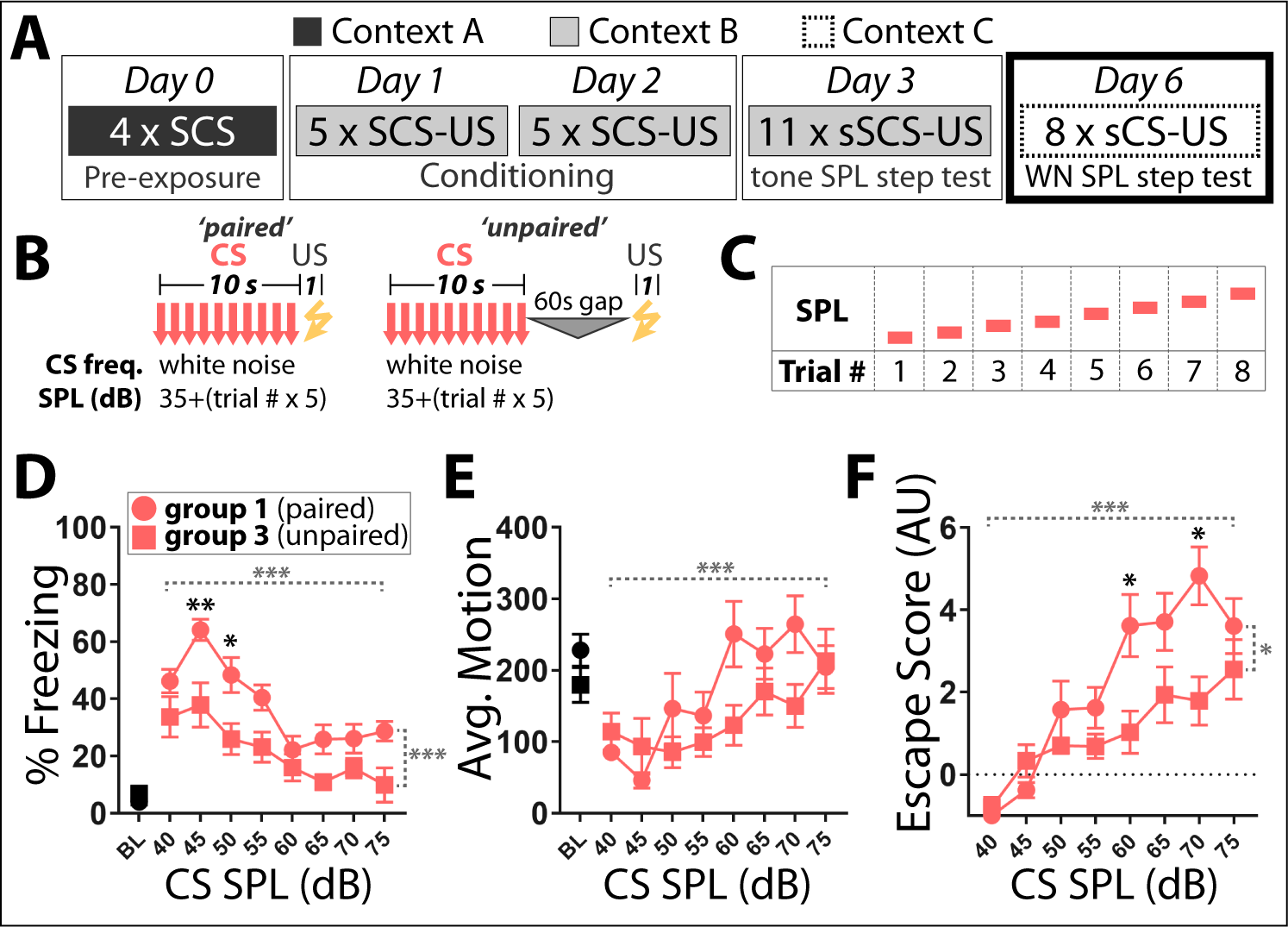
Sound pressure levels determine whether fear conditioned white noise stimuli elicit freezing or flight. (A) Mice conditioned in groups 1 and 3 (Figure 1) were run through a WN SPL step test in a novel context on day 6. (B) The WN step CS (sCS) is a white noise stimulus which begins at 40 dB and is stepped up by 5 dB each trial. (C) Schematic of WN SPL step test. (D) Percentage time spent freezing. (E) Average motion. (F) Escape score. Paired sCS, *n*=15; unpaired sCS, *n*=10. Black asterisks indicate significant difference between groups for a given trial. Dashed horizontal gray brackets indicates significant main effect of SPL. Dashed vertical gray brackets indicate significant main effect of group. Error bars indicate the SEM.

### 12 kHz tones trigger more conditioned escape behavior than 3 kHz tones in a pure tone SCS conditioning protocol

Elicitation of robust escape behavior by SCS conditioned 7.5 kHz tones required presentation at ≥90 dB (**Figure 2H**), whereas both paired and unpaired mice began responding actively to WN stimuli at SPL as low as 50 dB (**Figure 3D**). Although these stimuli differ in in terms of frequency, they also differ with regards to signal regularity: whereas the 7.5 kHz tone is sinusoidal and periodic, WN is random and aperiodic. Therefore, although the behavioral results detailed in Figure 1 could reflect differential sensitivity of mice to stimuli of different frequencies, they might alternatively be due to distinct defensive responses triggered by periodic versus aperiodic signals.

To directly test if frequency alone can influence defensive behaviors, we performed fear conditioning using a SCS composed of 3 and 12 kHz pure tones (**Figure 4A**). These frequencies were chosen as: *a)* the threshold SPL in mice is ∼100x lower for 12 kHz than 3 kHz pure tones [9]; perceived loudness of these two stimuli should thus differ when presented at standard SPL used during conditioning, similar to a 7.5 kHz/WN SCS; and *b)* 12 kHz is well separated from 17-20 kHz, a range that may be innately aversive in mice (see Discussion).

**Figure 4.**
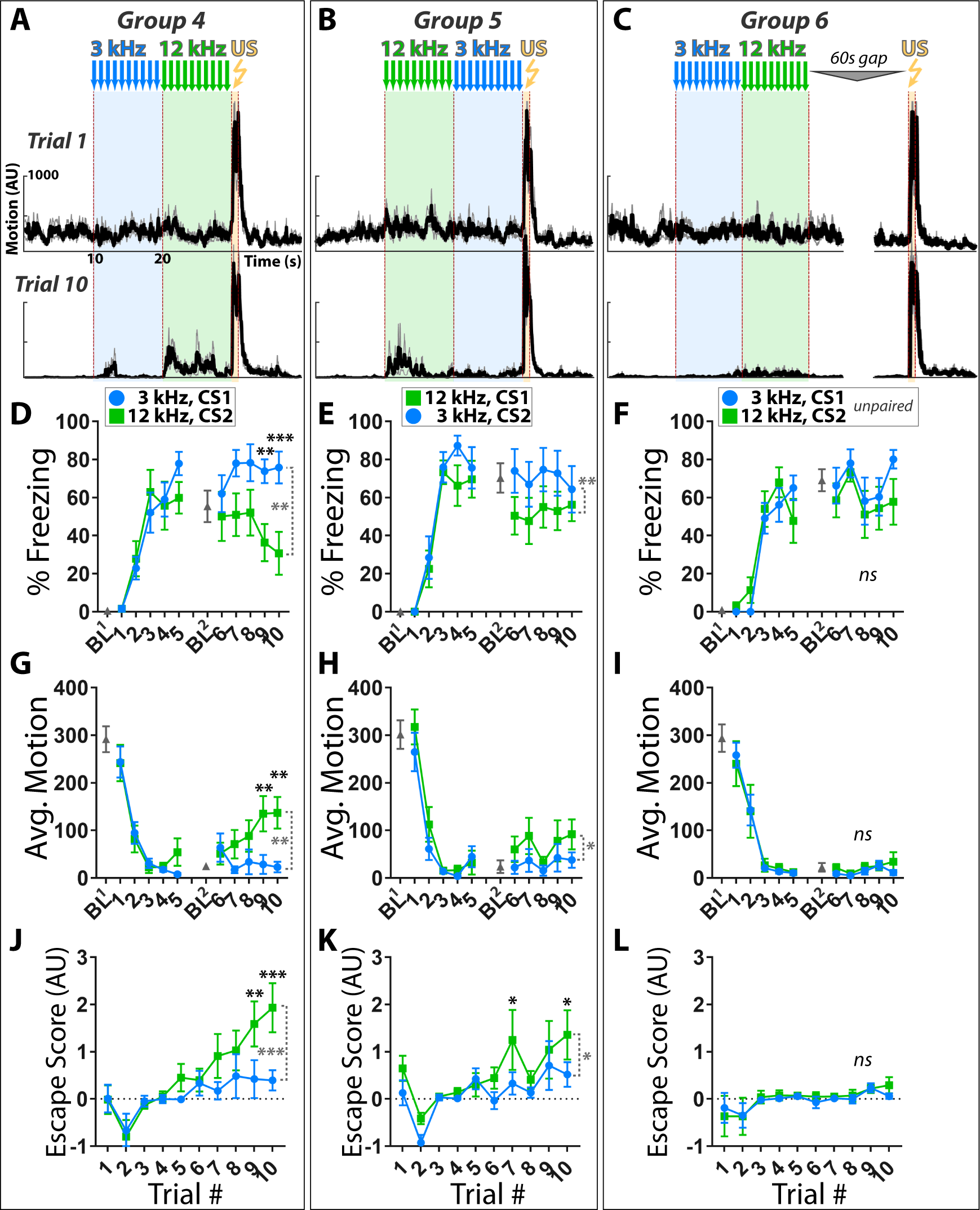
Active fear behaviors are more potently elicited by 12 kHz than 3 kHz stimuli during pure tone SCS conditioning. Contribution of CS audio frequency, order presented relative to US, and pairing were assessed by conditioning with a SCS composed of 3 and 12 kHz pure tones; conditioning done as in Figure 1A. (A-C) Motion indices (mean±SEM) show locomotor responses to stimuli (3 kHz pips, blue; 12 kHz pips, green; 0.9 mA footshock, yellow) for trials 1 (Day 1, *top*) and 10 (Day 2, *bottom*). (D-F) Percent time spent freezing. (G-I) Average motion. (J-L) Escape score. CS order and pairing: group 4 (A,D,G,J: CS1=3 kHz, CS2=12 kHz; *n*=10), group 5 (B,E,H,K: CS1=12 kHz, CS2=3 kHz; *n*=10), and group 3 (C,F,I,L: CS1=3 kHz, CS2=12 kHz, unpaired; *n*=10). Black asterisks indicate significant difference between groups for a given trial. Dashed vertical gray brackets indicate significant main effect of CS type. Error bars indicate the SEM.

As conditioning progressed, paired groups exhibited higher motion, less freezing, and more escape to the 12 kHz than 3 kHz CS, regardless of the order in which the stimuli were presented during training (**Figure 4,** groups 4 and 5). Notably, although mice in the unpaired group exhibited high levels of context fear, their behavioral responses to the two stimuli did not significantly differ, and no escape behaviors (darts or jumps) were observed by any mouse in this group in response to either stimulus during any trial (**Figures 4C,4F,4I,4L**). Importantly, the lack of active responses to the 3 kHz CS was not due to it being inaudible, as paired mice exhibited robust freezing to this stimulus when presented at an even lower SPL outside of the conditioning context (**Figure S2**). Thus, despite having no apparent intrinsic aversive valence, 12 kHz tones can elicit greater active fear responses than 3 kHz tones presented at equivalent SPL during SCS fear conditioning.

## DISCUSSION

In conclusion, we found that audio frequency properties strongly influence the defensive behaviors elicited by SCS fear conditioned auditory stimuli. Conditioned escape behaviors were most potently triggered by stimuli that contain frequencies to which mouse hearing is most sensitive, an effect that is independent of the order in which CSs were presented during learning. In addition, pure tones that elicit freezing at typical experimental sound pressure levels can promote conditioned escape when presented at higher levels. These data argue that stimulus salience, not temporal proximity to the US, is the primary means by which mice assess imminence and engage appropriate defensive strategies in the SCS paradigm.

Previous work provided compelling behavioral and neurophysiological evidence that SCS fear conditioned tone and white noise stimuli acutely elicit distinct defensive states indicative of different points along the threat imminence continuum [6]. We have demonstrated here that these defensive states track with the frequency and intensity of the conditioned stimuli, not order of CS presentation during learning. This indicates that mice are not shifting to more active threat coping strategies as perceived time to footshock decreases. Rather, these results argue that threat imminence in this model is determined primarily if not exclusively via the salience of threat-predictive auditory stimuli which, together with recent experience [10] and current fear levels, determine the threshold for switching from freezing to flight. This would appear to be similar mechanistically to how mice respond to innately threatening visual stimuli, where the probability and intensity of escape behaviors scale with the saliency of a threat stimulus [8]. Despite these similarities, it remains to be determined whether imminence and behavioral responses to threat stimuli of different sensory modalities are mediated via overlapping or distinct neural circuits. Indeed, separate pathways exist for processing fear of pain, predators, and social threats [11, 12], suggesting the possibility that visual and auditory fear stimuli might be processed via related yet discrete pathways. Consistent with this idea, distinct distributions of lateral amygdalar neurons were activated following fear conditioning with either an auditory or visual CS [13]. Recordings and functional manipulations of well-defined fear circuits in subjects sequentially exposed to visual (e.g. looming) and auditory (e.g. SCS) threat stimuli should help resolve this issue.

An implication of this work is the critical need to consider the behavioral sensitivity of experimental subjects to auditory stimuli of different frequencies. Psychophysical studies have demonstrated that all species have a particular range of frequencies that they hear well (i.e. which are audible at 10 dB); stimuli outside of this range may need to presented at substantially higher SPL in order to be efficiently detected. In addition, although most laboratory animals exhibit overlap in their hearing ranges, there can be significant differences in their sensitivity to particular frequencies, even among closely related species. For example, whereas the 10 dB threshold includes frequencies ranging from ∼5-40 kHz in rats, this range is very narrow in mice and limited to frequencies close to 16 kHz [14]. Differences can also exist across mouse strains and between different ages of the same strain. For example, C57BL/6J mice undergo hearing-loss induced plasticity that by 5 months of age results in loss of responsivity to high frequency tones (>20 kHz) with concomitantly enhanced behavioral sensitivity to middle (12 – 16 kHz) but not low (4 – 8 kHz) frequency stimuli [15, 16]. Moreover, certain frequencies may be innately aversive in rodents: rats emit and respond defensively to alarm vocalizations near 20 kHz [17-19], and 17-20 kHz ultrasonic sweeps can elicit robust freezing and flight behaviors in mice [8, 10]. White noise stimuli, which are both aperiodic and include 17-20 kHz frequencies, may thus be uniquely salient to mice under conditions of impending potential threats due to recruitment of dedicated defensive circuits tuned to innately threatening auditory stimuli. Consequently, discrimination studies that employ multiple auditory cues could be complicated both by variations in the ability of subjects to perceive different frequencies as well as potential innate valence associated with certain stimuli. For example, aversive conditioning to a 5 kHz CS+ followed by a generalization test using a higher salience CS-such as white noise could yield misleading conclusions if subjects exhibit escape behaviors to the CS- and, as is common, freezing is the only metric used to assess cue responsivity. Such confounds may be best avoided by assaying discrimination using tasks which measure behavioral responses to distinct patterns of a single, constant intensity sensory stimulus (e.g. drifting visual gratings of different orientation [20]). Interpretation of discrimination studies employing auditory stimuli would benefit from reversing the identities of the CS+ and CS-stimuli, and also from use of stimuli at frequencies and SPL that are detectable but do not trigger active fear behaviors.

Lastly, although reversing the order of the white noise and tone stimuli during training did not qualitatively alter the behaviors elicited by the CSs, this switch did have a quantitative effect. Specifically, white noise elicited significantly less conditioned escape behaviors when it preceded rather than followed the tone during training (Figure 1). One explanation for this result is that compound stimuli which increase in salience from CS1 to CS2 may produce greater arousal and learning than the reverse order. Indeed, tonal stimuli which sweep from low up to high frequencies are rated by human observers as more alarming than high to low sweeps [21]. Similarly, frequency upsweeps are associated with elevation of attention and arousal, whereas downsweeps are thought to have the opposite effect [22]. Use of compound stimuli that either increase or decrease in salience from CS1 to CS2 may thus have opposing influences on arousal in mice, an idea that could be tested in future studies by measuring arousal levels (e.g. using pupillometry) during training with different SCS conditioning protocols.

## METHODS

### Subjects

Male FVBB6 F1 hybrid mice, 4-5 months of age and weighing 25-30g were singly housed beginning one week prior to and throughout training and testing. All mice were maintained on a 12-hour reverse light/dark cycle with access to food and water *ad libitum*. The behavioral procedures used in this study were approved by the Institutional Animal Care and Use Committee at Boston Children’s Hospital.

### Apparatus

Behavioral training used fear conditioning chambers (30 X 25 X 25 cm, Med-Associates, Inc. St. Albans, VT), equipped with a Med-Associates VideoFreeze system. The boxes were enclosed in larger sound-attenuating chambers. Aspects of the boxes were varied to create two distinct contexts. The pre-exposure and testing context were composed of a white Plexiglas floor insert and a curved white Plexiglas wall insert with a hole over the wall speaker, making the rear walls of the chamber into a semi-circle. The ceiling and front door were composed of clear Plexiglas. The overhead light was off and the box was cleaned with 1% acetic acid. The conditioning context was comprised of a rectangular chamber with aluminum sidewalls and a white Plexiglas rear wall. The grid floor consisted of 16 stainless steel rods (4.8 mm thick) spaced 1.6 cm apart (center to center). Pans underlying each box were sprayed and cleaned between mice. Fans mounted above each chamber provided background noise (65dB). The experimental room was brightly lit with an overhead white light. Animals were kept in a holding room and individually transported to the experimental room in their home cage. Chambers were cleaned with soap and water following each day of behavioral testing.

### Serial Compound Stimulus (SCS) Fear Conditioning

For tone-white noise SCS, three groups of mice were conditioned with compound stimuli consisting of ten pure tone pips (7.5 KHz, 75 dB, 0.5s duration at 1Hz), ten white noise pips (WN, 75 dB, 0.5s duration at 1Hz), and a foot shock (0.9mA, 1s duration). The order and pairing differed between groups: Group 1 received Tone-WN paired with shock, Group 2 received WN-Tone paired with shock, and Group 3 received Tone-WN unpaired with shock (i.e. 60s gap in between CS2 and US). All groups had a 3 minute baseline period prior to the first CS and 30s after the final shock. Groups 1 and 2 had a 60s average pseudorandom ITI (range 50–90s), while Group 3 had a 180s average pseudorandom ITI (range 150-200). For pure tone SCS conditioning, the protocols were the same except that the tone and white noise stimuli were replaced with two pure tone stimuli: 3KHz (75dB, 10x 0.5s duration pips at 1Hz) and 12KHz (75dB, 10x 0.5s duration pips at 1Hz). On the day 0 of both experiments, mice were placed into the pre-exposure context and received four CS-alone trials. On Days 1 and 2, mice were placed into the conditioning context, where they received five CS trials that included shock. SPL step tests were run as indicated in the figures.

### Quantification of Behavior

Freezing behavior, average motion, and maximum motion were calculated using motion indices determined using automated near infrared (NIR) video tracking equipment and computer software (VideoFreeze, Med-Associates Inc.), as previously described [23]. Escape behaviors were scored manually from video files to count the number of darts and jumps. Darts were defined as rapid crossings preceded by immobility; jumps were defined as rapid movements in which all four paws left the floor. These behaviors were summed to determine the number of escape behaviors per mouse per trial, and used to quantify the vigor of responses to particular auditory stimuli via an ‘escape score’. As most mice were freezing (i.e. motion index = 0) throughout baseline periods on conditioning day 2, it was not possible to use a CS/BL motion index ratio as the basis of a ‘flight score’ as done previously [6]. Therefore, we calculated an ‘escape score’ ((MI_CS_ – MI_BL_)/100 + # of escape behaviors) by taking the difference in average motion index (MI) during CS versus the baseline for each trial (i.e. the 10 second period preceding delivery of a CS), dividing this by 100, and then adding 1 point for each dart or jump observed during that particular stimulus and trial.

### Statistical Analysis

Data were analyzed with paired t-tests or repeated-measures ANOVAs, with post hoc analysis correcting for multiple comparisons where appropriate. Statistical significance is labeled as *p<0.05, **p<0.01, and ***p<0.001.

## Supporting information

Video 1_crossing

Video 2_rapid movement in place

Video 3_Dart

Video 4_Jump

## ACKNOWLEDGEMENTS

We thank Delaney Foley for running the initial SCS conditioning experiments, and members of the Anthony lab for helpful discussions. This work was supported by NIH training grant #T32 NS007473 (S.H.), and grants from the Whitehall Foundation, Charles Hood Foundation, Tommy Fuss Center, Boston Children’s Hospital, Harvard Neurodiscovery Center, Harvard University Milton Fund, and Harvard Brain Initiative (T.E.A.).

---

**Supplemental Figure 1.**
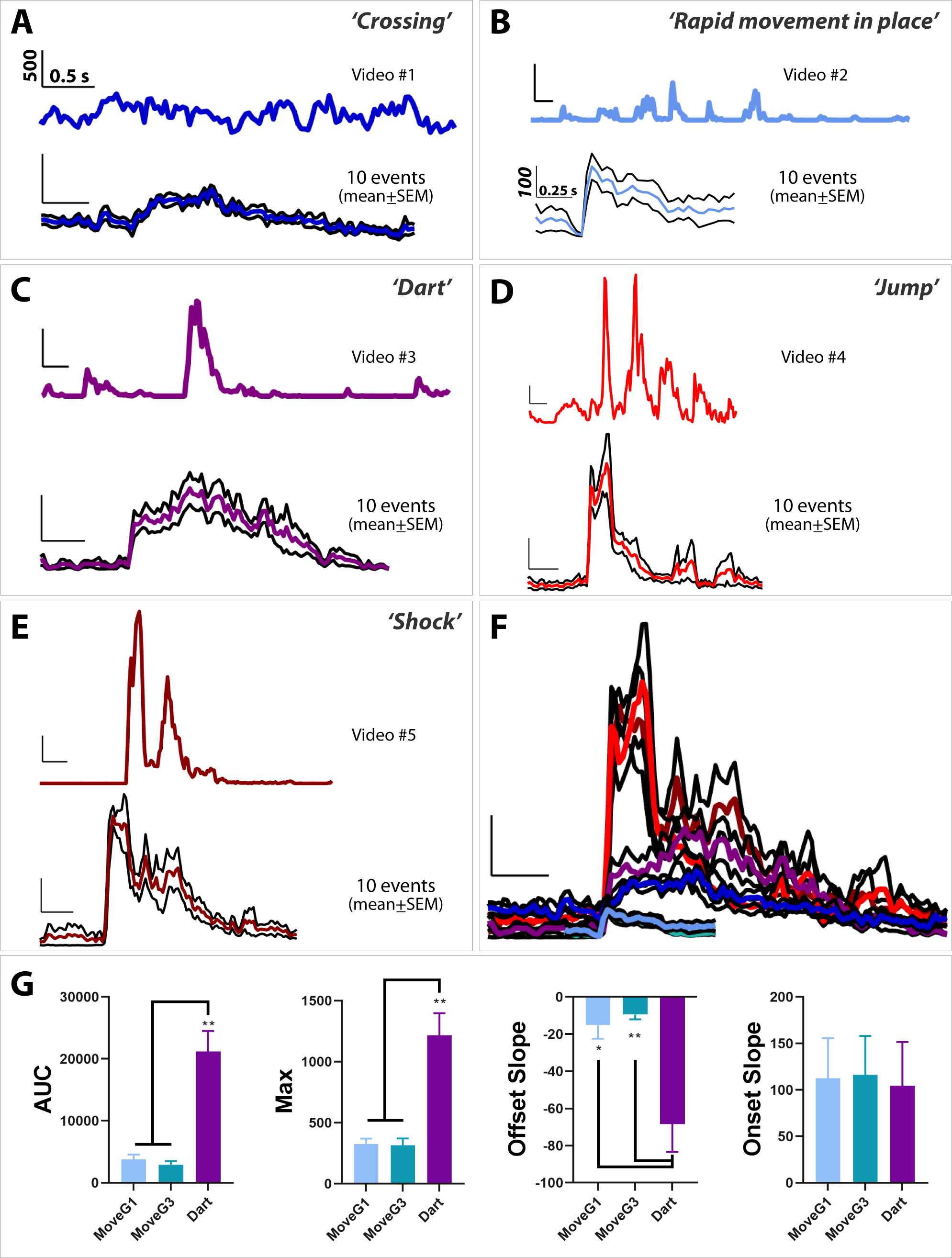
Manually identified behaviors exhibited during SCS conditioning can be quantitatively distinguished using motion index. A-F) Motion index traces of behaviors observed throughout serial compound stimulus fear conditioning. Upper traces show motion index across time for individual supplemental video examples. Lower traces show mean±SEM for 10 selected examples of each behavior. Scale bars: abscissa, time (sec)=0.25 (B), 0.5 (A,C-F); ordinate, motion index (AU)=100 (B), 500 (A,C-F). A) *Crossing* – which usually began on one wall of the chamber and ended on the opposite wall, were frequent prior to shock experience and decreased in frequency as mice exhibited freezing or other behaviors. Crossings that occurred during a CS presentation were not scored as escape behavior. B) ‘*Rapid movement in place’* – short, quick movements in a single location (usually in the corner) that include head movements (with or without wincing), turning in place, and shuffling feet without locomotion; these responses contributed to the time-locked motion responses to white noise pip onsets (Figure 1D), commonly occurring during WN presentations when escape behaviors were not observed, C) *Dart* – During early conditioning trials, darts were infrequent (Tone-WN experiment), or absent (3kHz-12kHz experiment), and increased in frequency and intensity throughout conditioning. These were eventually exhibited to CS stimuli by all mice in Groups 1 and 2, most mice in Group 3, most mice in Group 4, some mice in Group 5, and zero mice in Group 6. D) *Jump* – never occurred to CS stimuli in any group prior to SCS conditioning. On Day 2 of conditioning, these were exhibited to CS stimuli by the majority of mice in Group 1, a couple mice in Groups 2 and 3, and a subset of the mice in Groups 4 and 5. E) *Shock* – Mice typically responded to shock with one rapid-onset escape behavior and 2-4 subsequent escape behaviors; the most common profile was a jump followed by two darts. F) *Behavioral trace summary*. Overlay of average behavioral response for rapid movement in place (Group 1, light blue, Group 3, teal), Crossing (blue), Dart (purple), Jump (red), and Shock (dark red). G) *Comparison of WN-elicited behaviors*. The major behaviors that contributed to the time-locked motion responses to white noise were ‘rapid movements in place’ and darts. To quantitatively distinguish these two behaviors, ‘Rapid movements in place’ exhibited by mice in groups 1 (MoveG1) and 3 (MoveG3) were compared with ‘darts’ across several metrics (*n*=10 examples of each behavior per group). Maximum and area-under-the-curve (AUC) were computed for the duration of the observed behavior (until the behavior finished or another behavior emerged). To compute onset slope, a line was fitted from the beginning of the behavior to the max activity; for offset, a line was fitted from the max activity to the end of the behavior. As the variance was significantly different between groups, Welch ANOVA tests were run. Comparisons (Holm-Sidak) were made between all groups. Darts, in comparison to movements in place, had higher AUC (W(2,15.68)=13.88, p<0.001), higher maximum (W(2,16.11)=11.45, p<0.001), and a more negative offset slope (W(2,13.61)=7.37, p<0.01), but had the same onset slope (p>0.05), consistent with the rapid onset of the movement in place to each white noise onset.

**Supplemental Figure 2.**
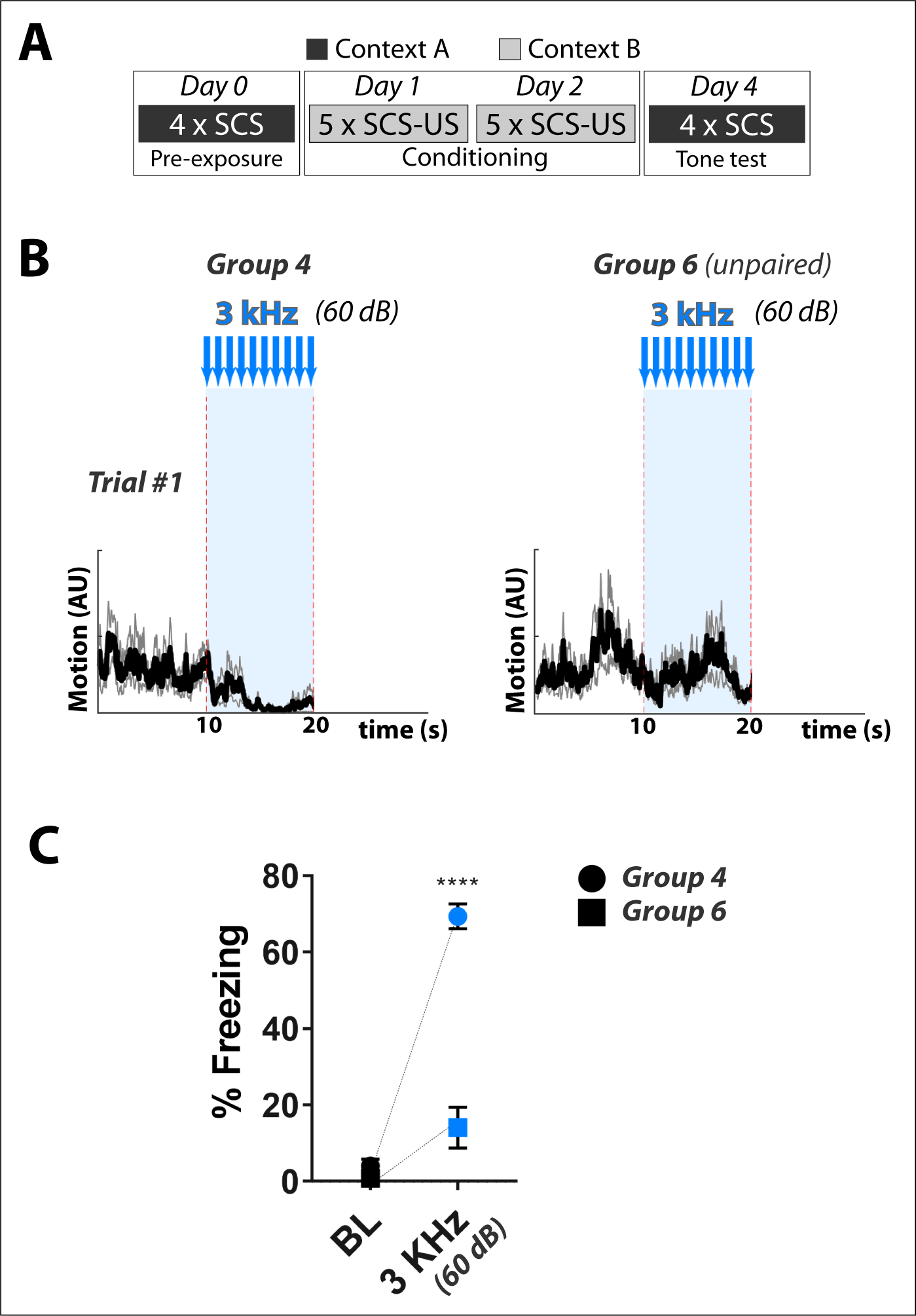
Mice hear and exhibit robust freezing to 3 kHz tone stimuli. (A) Tone tests were performed on mice conditioned using paired or unpaired SCS composed of 3 kHz (CS1) – 12 kHz (CS2) (Figure 4, main text). (B) Motion traces (mean+SEM) showing the first trial of the tone test in which 3 kHz tone pips at 60 dB were presented; note this is 15 dB lower than the SPL used during conditioning. (C) Mice in the paired group show robust freezing in response to tone presentation (paired t-test, freezing during 10 second baseline vs. freezing during 10 second 3 kHz tone presentation, p<0.0001) demonstrating that they can hear 3 kHz tone stimuli presented at the SPL used in the SCS experiments.

## REFERENCES

1. Blanchard, R.J., and Blanchard, D.C. (1989). Antipredator defensive behaviors in a visible burrow system. Journal of comparative psychology 103, 70–82.

2. Blanchard, R.J., Parmigiani, S., Bjornson, C., Masuda, C., Weiss, S.M., and Blanchard, D.C. (1995). Antipredator Behavior of Swiss-Webster Mice in a Visible Burrow System. Aggressive Behavior 21, 123–136.

3. Bouton, M.E., and Bolles, R.C. (1980). Conditioned fear assessed by freezing and by the suppression of three different baselines. Animal Learning & Behavior 8, 429–434.

4. Fanselow, M.S. (1994). Neural organization of the defensive behavior system responsible for fear. Psychonomic bulletin & review 1, 429–438.

5. Fanselow, M.S., and Lester, L.S. (1988). A functional behavioristic approach to aversively motivated behavior: predatory imminence as a determinant of the topography of defensive behavior. In Evolution and learning, R.C. Bolles and M.D. Beecher, eds. (Hillsdale, NJ: Erlbaum), pp. 185 – 211.

6. Fadok, J.P., Krabbe, S., Markovic, M., Courtin, J., Xu, C., Massi, L., Botta, P., Bylund, K., Muller, C., Kovacevic, A., et al. (2017). A competitive inhibitory circuit for selection of active and passive fear responses. Nature 542, 96–100.

7. Yilmaz, M., and Meister, M. (2013). Rapid innate defensive responses of mice to looming visual stimuli. Current biology : CB 23, 2011–2015.

8. Evans, D.A., Stempel, A.V., Vale, R., Ruehle, S., Lefler, Y., and Branco, T. (2018). A synaptic threshold mechanism for computing escape decisions. Nature 558, 590–594.

9. Koay, G., Heffner, R., and Heffner, H. (2002). Behavioral audiograms of homozygous med(J) mutant mice with sodium channel deficiency and unaffected controls. Hearing research 171, 111–118.

10. Mongeau, R., Miller, G.A., Chiang, E., and Anderson, D.J. (2003). Neural correlates of competing fear behaviors evoked by an innately aversive stimulus. The Journal of neuroscience : the official journal of the Society for Neuroscience 23, 3855–3868.

11. Gross, C.T., and Canteras, N.S. (2012). The many paths to fear. Nature reviews. Neuroscience 13, 651–658.

12. Silva, B.A., Mattucci, C., Krzywkowski, P., Murana, E., Illarionova, A., Grinevich, V., Canteras, N.S., Ragozzino, D., and Gross, C.T. (2013). Independent hypothalamic circuits for social and predator fear. Nature neuroscience 16, 1731– 1733.

13. Bergstrom, H.C., and Johnson, L.R. (2014). An organization of visual and auditory fear conditioning in the lateral amygdala. Neurobiology of learning and memory 116, 1–13.

14. Heffner, H.E., and Heffner, R.S. (2007). Hearing ranges of laboratory animals. Journal of the American Association for Laboratory Animal Science : JAALAS 46, 20–22.

15. Carlson, S., and Willott, J.F. (1996). The behavioral salience of tones as indicated by prepulse inhibition of the startle response: relationship to hearing loss and central neural plasticity in C57BL/6J mice. Hearing research 99, 168– 175.

16. Willott, J.F., Carlson, S., and Chen, H. (1994). Prepulse inhibition of the startle response in mice: relationship to hearing loss and auditory system plasticity. Behavioral neuroscience 108, 703–713.

17. Beckett, S.R., Aspley, S., Graham, M., and Marsden, C.A. (1996). Pharmacological manipulation of ultrasound induced defence behaviour in the rat. Psychopharmacology 127, 384–390.

18. Blanchard, R.J., Agullana, R., McGee, L., Weiss, S., and Blanchard, D.C. (1992). Sex differences in the incidence and sonographic characteristics of antipredator ultrasonic cries in the laboratory rat (Rattus norvegicus). Journal of comparative psychology 106, 270–277.

19. Cuomo, V., Cagiano, R., De Salvia, M.A., Mazzoccoli, M., Persichella, M., and Renna, G. (1992). Ultrasonic vocalization as an indicator of emotional state during active avoidance learning in rats. Life sciences 50, 1049–1055.

20. Burgess, C.R., Ramesh, R.N., Sugden, A.U., Levandowski, K.M., Minnig, M.A., Fenselau, H., Lowell, B.B., and Andermann, M.L. (2016). Hunger-Dependent Enhancement of Food Cue Responses in Mouse Postrhinal Cortex and Lateral Amygdala. Neuron 91, 1154–1169.

21. Catchpole, K.R., McKeown, J.D., and Withington, D.J. (2004). Localizable auditory warning pulses. Ergonomics 47, 748–771.

22. Owren, M.J., and Rendall, D. (2001). Sound on the Rebound: Bringing form and function back to the forefront in understanding nonhuman primate vocal signaling. Evolutionary Anthropology 10, 58–71.

23. Zelikowsky, M., Bissiere, S., Hast, T.A., Bennett, R.Z., Abdipranoto, A., Vissel, B., and Fanselow, M.S. (2013). Prefrontal microcircuit underlies contextual learning after hippocampal loss. Proceedings of the National Academy of Sciences of the United States of America 110, 9938–9943.

